# *Chromulinavorax destructans*, a pathogenic TM6 bacterium with an unusual replication strategy targeting protist mitochondrion

**DOI:** 10.1101/379388

**Authors:** Christoph M. Deeg, Matthias M. Zimmer, Emma George, Filip Husnik, Patrick J. Keeling, Curtis A. Suttle

## Abstract

Most of the diversity of microbial life is not available in culture, and as such we lack even a fundamental understanding of the biological diversity of several branches on the tree of life. One branch that is highly underrepresented is the candidate phylum TM6, also known as the Dependentiae. Their biology is known only from reduced genomes recovered from metagenomes around the world and two isolates infecting amoebae, all suggest that they live highly host-associated lifestyles as parasites or symbionts. *Chromulinavorax destructans* is an isolate from the TM6/Dependentiae that infects and lyses the abundant heterotrophic flagellate, *Spumella elongata. Chromulinavorax destructans* is characterized by a high degree of reduction and specialization for infection, so much so it was discovered in a screen for giant viruses. Its 1.2 Mb genome shows no metabolic potential and *C. destructans* instead relies on extensive transporter system to import nutrients, and even energy in the form of ATP from the host. Accordingly, it replicates in a viral-like fashion, while extensively reorganizing and expanding the host mitochondrion. 44% of proteins contain signal sequences for secretion, which includes many proteins of unknown function as well as 98 copies of ankyrin-repeat domain proteins, known effectors of host modulation, suggesting the presence of an extensive host-manipulation apparatus.

## Introduction

The vast majority of genetic diversity on earth is encoded by microorganisms, most of which are not represented in culture and are only known from barcoding or metagenomic studies [1]. As a result, there are many major branches in the tree of life where we know little or sometimes nothing at all about the biological diversity represented by the tree. One such case is the candidate phylum TM6, also known as Dependentiae, which was originally described from a 16S environmental amplicon library prepared from a peat sample collected in northern Germany [2]. Later, low complexity metagenomic data from a hospital biofilm, soil, and waste water allowed for the assembly of almost complete genomes from representatives of the TM6 phylum [3, 4]. These genomes implied very limited metabolic capability, while encoding an extensive system of transporters, including ATP transporters, suggesting that TM6 bacteria may be symbionts or parasites that rely extensively on their hosts for energy and metabolites [3, 4]. The genomic complement, enriched in conserved gene functions found in endosymbionts or parasites of eukaryotes such as *Chlamydia, Wolbachia*, and *Rickettsia*, further suggested that TM6 was associated with eukaryotic hosts [3, 4]. Indeed, two representatives of the TM6 phylum were since isolated in amoebae, competent hosts to a vast array of bacterial, eukaryotic, and viral symbionts and pathogens [5, 6]. The first isolate, *Babela massiliensis*, is an obligate pathogen that causes lysis of its host, *Acanthamoeba castellanii*, while the other, *Vermiphilus pyriformis*, maintains a stable relationship with its host *Vermamoeba vermiformis*, suggesting that different TM6 bacteria might employ different life history strategies [5, 6]. Since only the sequence for *Babela massiliensis* is available, a detailed comparison of the genomic complement involved in the different lifestyles is not possible. Both isolates show unusual replication strategies within their hosts that delay cell fission and initially produce large replication bodies before dividing into progeny cells. Recent advances in metagenomics and single-cell genomics have added further almost-complete metagenomically assembled genomes (MAGs) to the proposed candidate phylum that seem to be distinct from the amoeba-infecting species; however, the interpretation of their genomic complement remains speculative with only amoeba-infecting isolates reported to be in culture [7–9].

Amoebae are probably the best studied microbial eukaryotic hosts of bacterial symbionts and pathogens, but represent only a fraction of eukaryotic diversity. Another large and polyphonic group of microbial eukaryotes that are abundant in most natural environments are heterotrophic nanoflagellates, defined as being between 2 and 20 μm in size, and by their lifestyle of preying on bacteria, viruses, and other microbial eukaryotes [10, 11]. They are important in aquatic ecosystems, as the component of the microbial loop that most directly links to higher trophic levels. Grazing by nanoflagellates, together with viral lysis, are the primary mortality agents of microbial populations in aquatic environments [12]. Chrysophytes are a diverse group of nanoflagellates within the stramenopiles, and include a number of mixotrophs in the genera *Ochromonas* and *Chromulina*, for example, as well as pure heterotrophs, such as members of the genus *Spumella* [13, 14]. Present in fresh and salt waters, *Spumella* is one of Earth’s most abundant phagotrophic nanoflagellates and is readily isolated from many environments [15].

Here, *Chromulinavorax destructans*, a novel isolate of the TM6/Dependentiae, is described which infects and lyses *Spumella elongata*. Its unusual replication cycle superficially resembles the infection cycle of a virus, and includes the creation of a replication ‘factory’ and the remodelling of the host mitochondrion within its host cell. The genome is highly reduced and lacks nearly all genes related to energy metabolism. Moreover, like MAGs of other TM6, it encodes a large suite of transporters, including ATP transporters. Overall, *C. destructans* provides a link between molecular survey data and biological observations, and expands the known functional diversity and ecological roles of TM6 bacteria.

## Materials and methods

### Sampling

Samples were collected from 11 freshwater locations in southern British Columbia, Canada (49°49’4”N, 123° 7’46”W; 49°42’5”N, 123° 8’47”W; 49°37’34”N, 123°12’27”W; 49° 6’12”N, 122° 4’38”W; 49° 5’22”N, 122° 7’1”W; 49°18’10”N, 122°42’9”W; 49° 8’27”N; 123° 3’16”W; 49°13’21”N, 123°12’43”W; 49°13’13”N, 123°12’41”W; 49°14’52”N, 123°13’59”W; 49°15’58”N, 123°15’34”W. See Chapter 2). To concentrate pathogens, 20 liter water samples were prefiltered with a GF-A filter (Millipore, Bedford, MA, USA; nominal pore size 1.1 μm) over a 0.8-μm pore-size PES membrane (Sterlitech, Kent, WA, USA) [16]. Filtrates from all locations were pooled and concentrated using a 30kDa MW cut-off tangential flow ultrafiltration cartridge (Millipore, Bedford, MA, USA) [17].

### Isolation

The host organism, *Spumella elongata* strain CCAP 955/1 was kindly provided by David Caron (University of Southern California) and maintained in modified DY-V artificial fresh water media with yeast extract and a wheat grain [18]. Cultures of *S. elongata* at approximately 2×10^5^ cells/ml were inoculated with the pooled microbial concentrates from all 11 locations. Cell numbers of the inoculated culture were monitored by flow cytometry and compared to a medium-only mock-infected control culture using flow cytometry (LysoTracker Green (Molecular Probes) vs. FSC on FACScalibur (Becton-Dickinson, Franklin Lakes, New Jersey, USA)) [19]. After cell lysis, the lysate was filtered through a 0.8-μm pore-size PES membrane (Sterlitech) to remove remaining host cells. The lytic agent was propagated and made clonal by three serial end-point dilutions. The concentrations of the lytic agent were screened by flow cytometry using SYBR Green (Invitrogen Carlsbad, California, USA) nucleic-acid stain after 2% glutaraldehyde fixation (vs SSC). The flow cytometry profile presented as a population clearly distinct from heterotrophic bacteria, phage, and eukaryotes; however, their larger size heterogeneity when compared to giant virus isolates suggested a non-viral nature [20].

### Transmission electron microscopy

#### Negative staining

Lysates of *Spumella elongata* after infection with *Chromulinavorax destructans* were applied to the carbon side of a formvar carbon-coated 400-mesh copper grids (TedPella, CA, USA) and incubated at 4°C in the dark overnight in the presence of high humidity. Next, the lysate was removed and the grids were stained with 1% Uranyl acetate for 30 s.

#### Ultra-thin sectioning

Exponentially growing cultures of *S. elongata* at a concentration of 5×10^5^ cells ml^−1^ were infected with *C. destructans* at a ratio of ~5 pathogen to host cells to ensure synchronous infection. Cells were harvested from infected cultures at 3, 6, 9, 12, 18, and 24 h post infection, as well as from uninfected control cultures. Cells from 50 ml were pelleted in two consecutive 10 min at 5000 xg centrifugation runs in a fixed angle Beckmann tabletop centrifuge. For chemical fixation, the pellet was resuspended in 0.2 M Na cacodylate buffer, 0.2 M sucrose, 5% EM grade glutaraldehyde, pH 7.4 and incubated for 2 h on ice. After washing in 0.2 M Na cacodylate buffer, cells were postfixed with 1% osmium tetroxide. Samples were dehydrated through water/ethanol gradients and ethanol was substituted by acetone. An equal part mixture of Spurr’s and Gembed resin was used to embed the cells, which was polymerized at 60°C overnight.

For high pressure freezing, cell pellets were resuspended in 10-15 μl of DY-V culture medium with 20% (w/v) BSA and immediately placed on ice. Cell suspensions were cryo-preserved using a Leica EM HPM100 high-pressure freezer. Vitrified samples were freeze-substituted in a Leica AFS system for 2 d at −85°C in a 0.5% glutaraldehyde / 0.1% tannic acid solution in acetone, then rinsed ten times in 100% acetone at −85°C, and transferred to 1% osmium tetroxide, 0.1% uranyl acetate in acetone and stored for an additional 2 d at −85°C. The samples were then warmed to −20°C for 10 h, held at −20°C for 6 h to facilitate osmication, and then warmed to 4°C for 12 h. The samples were then rinsed in 100% acetone 3X at room temperature and gradually infiltrated with an equal part mixture of Spurr’s and Gembed embedding media. Samples were polymerized in a 60°C oven overnight. Fifty nm thin sections were prepared using a Diatome ultra 45° knife (Diatome, Switzerland) on an ultra-microtome. The sections were collected on a 400x copper grid and stained for 10 min in 2% aqueous uranyl acetate and 5 min in Reynold’s lead citrate. Image data were recorded on a Hitachi H7600 transmission electron microscope at 80 kV. Image J (RRID:SCR_003070) was used to compile all TEM images. Adjustments to contrast and brightness levels were applied equally to all parts of the image.

### Pathogen concentration and sequencing

For PacBio sequencing, exponentially growing *S. elongata* cultures at a concentration of approximately 5×10^5^ cells ml^−1^ were infected with *C. destructans* lysate (~10^7^ cells ml^−1^) at a multiplicity of infection (MOI) of ~0.5. After four days, when host cells were undetectable, cultures were centrifuged in a Sorvall SLC-6000 centrifuge with fixed angle rotor for 20 min and 5000 rpm at 4°C to remove remaining host cells and the supernatant was subjected to tangential flow filtration with at 30kDa cut-off (Vivaflow PES) and concentrated approximately 100x. Concentrates were ultracentrifuged at 28,000 rpm, 15°C for 8h in a Ti90 fixed-angle rotor (Beckman-Coulter, Brea, California, USA). Next, the concentrate was further concentrated by sedimenting it onto a 40% Optiprep 50 mM Tris-Cl, pH 8.0, 2mM MgCl_2_ cushion for 30 min at 28,000 rpm and 15°C in a SW40Ti swinging-bucket rotor in an ultracentrifuge (Beckman-Coulter, Brea, California, USA). An Optiprep (Sigma) gradient was created by underlaying a 10% Optiprep solution in 50 mM Tris-Cl, pH 8.0, 2 mM MgCl_2_ with a 30% Optiprep solution followed by a 50% Optoprep solution and was equilibration overnight at 4°C. One ml of concentrate from the 40% cushion was added atop the gradient and the concentrate was fractionated by ultracentrifugation in an SW40 rotor for 4 h at 25000 rpm and 18°C. The fraction corresponding to the pathogen was extracted from the gradient with a syringe and washed twice with 50 mM Tris-Cl, pH 8.0, 2 mM MgCl_2_ followed by centrifugation in an SW40 rotor for 20 min at 7200 rpm and 18°C and finally collected by centrifugation in an SW40 rotor for 30 min at 7800 rpm and 18°C. Purity of the concentrate was verified by flow cytometry (SYBR Green (Invitrogen Carlsbad, California, USA) vs SSC on a FACScalibur flow cytometer (Becton-Dickinson, Franklin Lakes, New Jersey, USA). High molecular weight genomic DNA was extracted using phenol-chloroform-chloroform extraction. Length and purity of the DNA were confirmed by gel electrophoresis and a Bioanalyzer 2100 with the HS DNA kit (Agilent Technology). PacBio RSII 20kb sequencing was performed by the sequencing center of the University of Delaware. Reads were assembled using PacBio HGAP3 software with 20 kb seed reads and resulted in a single contig of 1,228,924bp, 819.19x coverage, 100% called bases and a consensus concordance of 99.9839% [21].

### Annotation

The genome was circularized, resulting in a final chromosome size of 1,174,272 bp. Genome annotation was performed using the automated NCBI Prokaryotic Genome Annotation Pipeline (PGAAP). In parallel, open reading frames were predicted using GLIMMER (RRID:SCR_011931) with default settings [22]. Translated proteins were analyzed using BLASTp, CDD RPS-BLAST and pfam HMMER. These results were used to refine the PGAAP annotation. Signal peptides and trans-membrane domains were predicted using Phobius [23]. The annotated genome is available under the accession number CP025544. Metabolic pathways were predicted using the Kyoto Encyclopedia of Genes and Genomes (KEGG RRID:SCR_012773) automatic annotation server KAAS and Pathway Tools (RRID:SCR_013786) [24, 25].

### Phylogenetic analysis

Full length 16S rDNA sequences belonging to the candidate phylum TM6 were downloaded from NCBI. Alignments of rDNA sequences were done in Geneious R9 (RRID:SCR_010519) using MUSCLE with default parameters (RRID:SCR_011812)[26]. Maximum likelihood trees were constructed with RAxML ML search with 1000 rapid bootstraps using GTR+GAMMA [27].

Near-complete metagenomically assembled genomes (MAGs) and complete genomes of isolates were retrieved from NCBI. There were 3820 clusters of orthologous genes defined by OrthoMCL (RRID:SCR_007839) with standard parameters (Blast E-value cut-off = 10^−5^ and mcl inflation factor = 1.5) on all protein coding genes of length ≥ 100 aa [28]. Overlap in gene content was defined using a custom R script (see Deeg et al. 2018 [20]). Genes encoding ribosomal proteins were identified by BLAST. Ribosomal protein L2, L3, L4, L5, L6, L14, L15, L18, L16, L22, L24, S3, S8, S10, S17, and S19 sequences were aligned using MUSCLE with default parameters (RRID:SCR_011812) [26]. Maximum-likelihood trees were constructed with RAxML rapid bootstrapping and ML search with 1000 bootstraps utilizing GTR+CAT [27].

## Results

### Chromulinavorax destructans is a lytic pathogen of Spumella elongata

In an effort to isolate pathogens that infect ecologically relevant protistan zooplankton, microbial assemblages smaller than 0.8 μm from a variety of freshwater habitats in southwestern British Columbia were concentrated (Chapter 2). The concentrates were used to inoculate cultures of protists isolated from the same environments, as well as from culture collections, and cell densities were monitored by flow cytometry. One such screen yielded a pathogen infecting *Spumella elongata* (CCAP 955/1) that caused complete lysis of the culture within 48 h post infection (hpi). The lytic agent was named *Chromulinavorax destructans* (subsequently referred to as *C. destructans)*. Infection was strain-specific and a different isolate of *Spumella sp*. from one of the environments sampled for pathogen concentration did not cause lysis. The pathogen continued to cause lysis of *S. elongata* after storage at 4°C for up to four years. Further, treatment with a bacteriocidal multi-antibiotic cocktail containing Ampicillin, Vancomycin, and Rifampicin also did not ablate the lytic potential. Under epifluorescence microscopy, infected cells developed large DNA-containing intracytoplasmic compartments starting at 8 hpi, followed by lysis starting at 18 hpi that was associated with numerous DNA-containing particles of approximately 0.4 μm bursting out of the cytoplasm (Figure 1B). *Chromulinavorax destructans* cells were first observed by flow cytometry 12 hpi (Figure 2 B), coinciding with a decline in hostcell density (Figure 2A). The population of *C. destructans* showed a homogeneous fluorescence signature suggesting that DNA replication does not occur outside of the host (Figure 2B). Side-scatter heterogeneity suggests that the cells vary in size, or are clumped (Figure 2B). In negative-staining electron microscopy, *C. destructans* cells present as 350-400 nm cocci with one polymorphous depression, similar to *Babela massiliensis* (Figure 1C, Figure 2C), [6]. In thin-section electron microscopy, free *C. destructans* cells show a gram-negative-like coccoid morphology with a lipid double layer and electron-dense material in the periplasm (Figure 1D). The central part of the cytoplasm shows an electron-dense nucleoid. No depression similar to that seen in negative staining was observed in thin sections, suggesting that this is a preparation artefact of negative staining.

**Figure 1:**
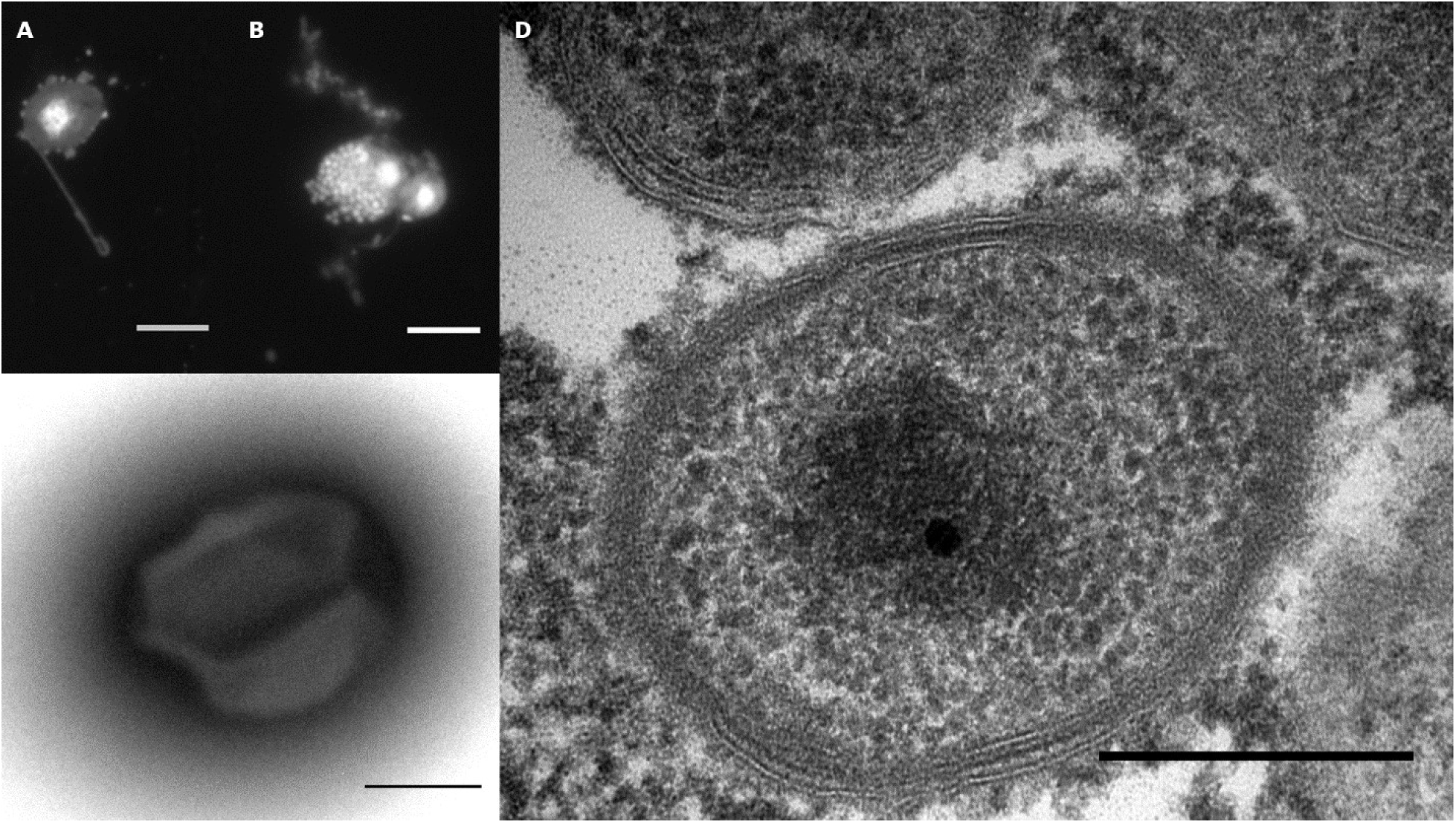
*Micrographs of* Chromulinavorax destructans. A: Epifluorescence micrograph of DAPI-stained *Spumella elongata* (prominently-stained nucleus). B: Epifluorescence micrograph of two *S. elongata* cells 19 h after exposure to *C. destructans*. Coccoid cells of *C. destructans* are seen bursting from a cell undergoing lysis. C: Purified *C. destructans* cell in negative-staining electron micrographs show a depression on the cell surface. D: Thin-sectioned, high-pressure frozen electron micrographs of *C. destructans* reveal 350- to 400-nm particles with inner and outer membranes, as well as dark-staining periplasmic material. The genome is condensed into a nucleoid during dormancy. Scale bars: A, B: 5 μm; C, D: 250 nm.

**Figure 2:**
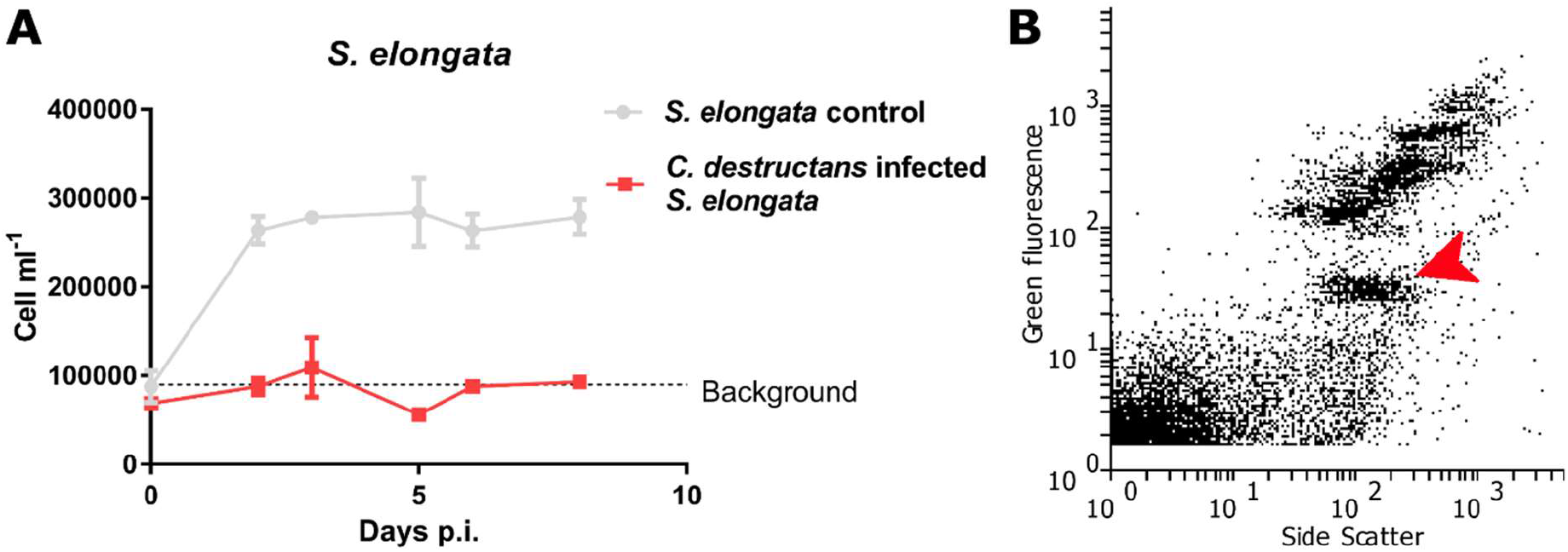
*Effect of addition of* Chromulinavorax destructans *to a culture of* Spumella elongata: A: Change in the cell number of *S. elongate* with (infected) and without (control) exposure *to C. destructans* is added (detection limit 10^5^ cells ml^−1^). The error bars represent the SEM of triplicate cultures, and the asterisks, time points where the differences between the control and infected cultures were significantly different (p<0.005). B: Flow cytometric profile at 48hpi of *C. destructans* (red arrow) in a culture of S. elongata and stained with SYBR-Green.

### *Chromulinavorax destructans* replicates in bodies nested in large mitochondrial invaginations

The life cycle of the lytic bacterium *Chromulinavorax destructans* putatively begins when ingested by the phagotrophic protist, *Spumella elongata*, a heterokont and slightly elongated flagellate (Figure 1A, Figure 3A, Figure 4C), although it can also form larger amoeboid cells. Once in the food vacuoles, the cells appear to secrete outer membrane vesicles (Figure 4A/B), and by 3 hpi they appear as a spherical mass in the host’s cytoplasm with the mitochondrion partially wrapped around the bacterial replication body by forming a deep invagination (Figure 3B). The invagination becomes more pronounced over time until the mitochondrion completely surrounds the replication body. During this process, the membrane integrity of the parasitoid and the mitochondrion, including its cristae, remain intact, suggesting that there is no invasion of the mitochondrial matrix (Figure 3C). At 9 hpi, the bacterial replication bodies are surrounded by the mitochondrion, and appear as several amorphous elongated bodies (Figure 4). This expansion phase culminates around 12 hpi, when the mitochondrion, now containing numerous invaginations and inclusions occupied by the replication bodies, takes up two thirds of the host cytoplasm (Figure 3D). Despite this extensive modification, the mitochondrion is still intact as membrane integrity is not compromised and cristae structures are preserved (Figure 3D). This integrity is contrasted by the degradation of the nucleus, as well as extensive membrane disarray inside the cell, including membrane vesicles budding from the cell (Figure 3D). The *C. destructans* replication bodies now show signs of regularly spaced invaginations, preceding division into the mature cocci (Figure 3D). The replication cycle completes at around 19 hpi, when *Spumella elongata* ghost cells with emptied cytoplasm appear (Figure 3E). The mature coccoid form of *C. destructans* are seen both intracellularly and extracellularly, presumably released by bursting of the compromised host cell membrane (Figure 3E, Figure 3E). Occasionally, replication bodies that did not complete the replication cycle, possibly due to prematurely exhausting the host cell’s resources, were observed within ghost cells (Figure 3E, Figure 3E).

**Figure 3:**
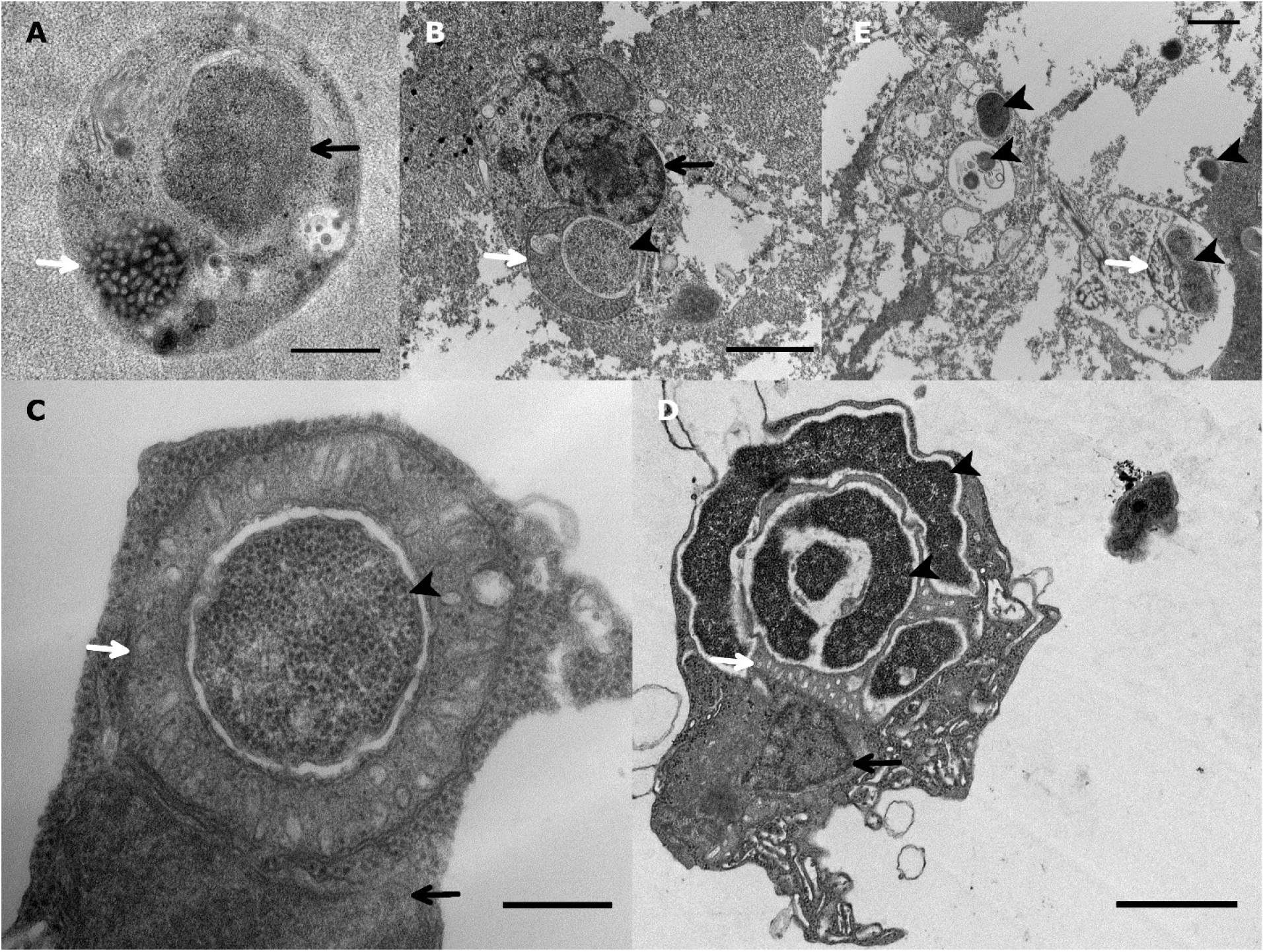
*Replication cycle of Chromulinavorax destructans*. A: Electron micrograph of a healthy *Spumella elongata* cell showing the nucleus (black arrow) and the mitochondrion (white arrow). B: At 3 hpi, *C. destructans* (black arrow head) has invaded the host cytoplasm and the mitochondrion (white arrow) has wrapped around the replication body, while the nucleus stays intact (black arrow). C: Close-up of the replication body (black arrow head) surrounded by the host mitochondrion (white arrow) showing intact membranes of both the pathogen and the organelle (black arrow: nucleus). D: At 12 hpi, *C. destructans* shows long filaments and the beginning of septation (black arrowhead), surrounded by the highly invaginated mitochondrion (white arrow). The nucleus is present, but shows signs of degradation. E: Ghost cells of *S. elongata* with degraded cytoplasm and organelles (white arrow: mitochondrion) and *C. destructans* (black arrow heads) at 18 hpi. Scale bars: A: 500 nm; C: 250 nm; B,D,E: 1 μm

**Figure 4:**
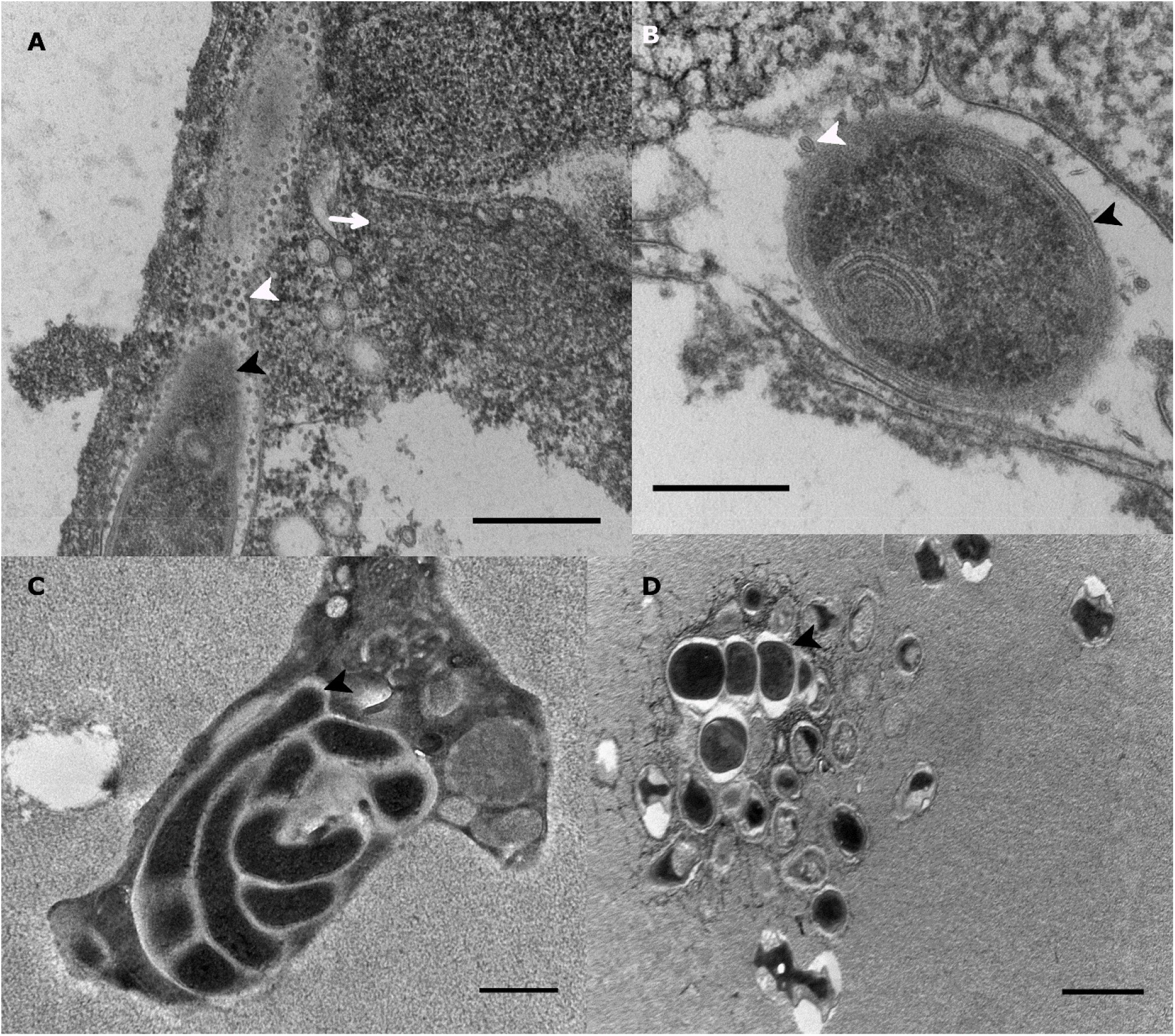
*Replication cycle of* Chromulinavorax destructans. A: Putative *C. destructans* (black arrow head) inside the food vacuole of *Spumella elongata* showing apparent secretion of outer membrane vesicles (white arrow head) and an intact S. elongata mitochondrion (white arrow). B: Putative *C. destructans* (black arrow head) inside the food vacuole of *Spumella elongata* showing liposomes and outer membrane vesicles (white arrow head). C: Extensive *C. destructans* replication inside *S. elongata* at 9 hpi. D: Mature cells of *C. destructans* being released from a ghost cell of *S. elongata* at 24 h post-infection.

**Figure 5:**
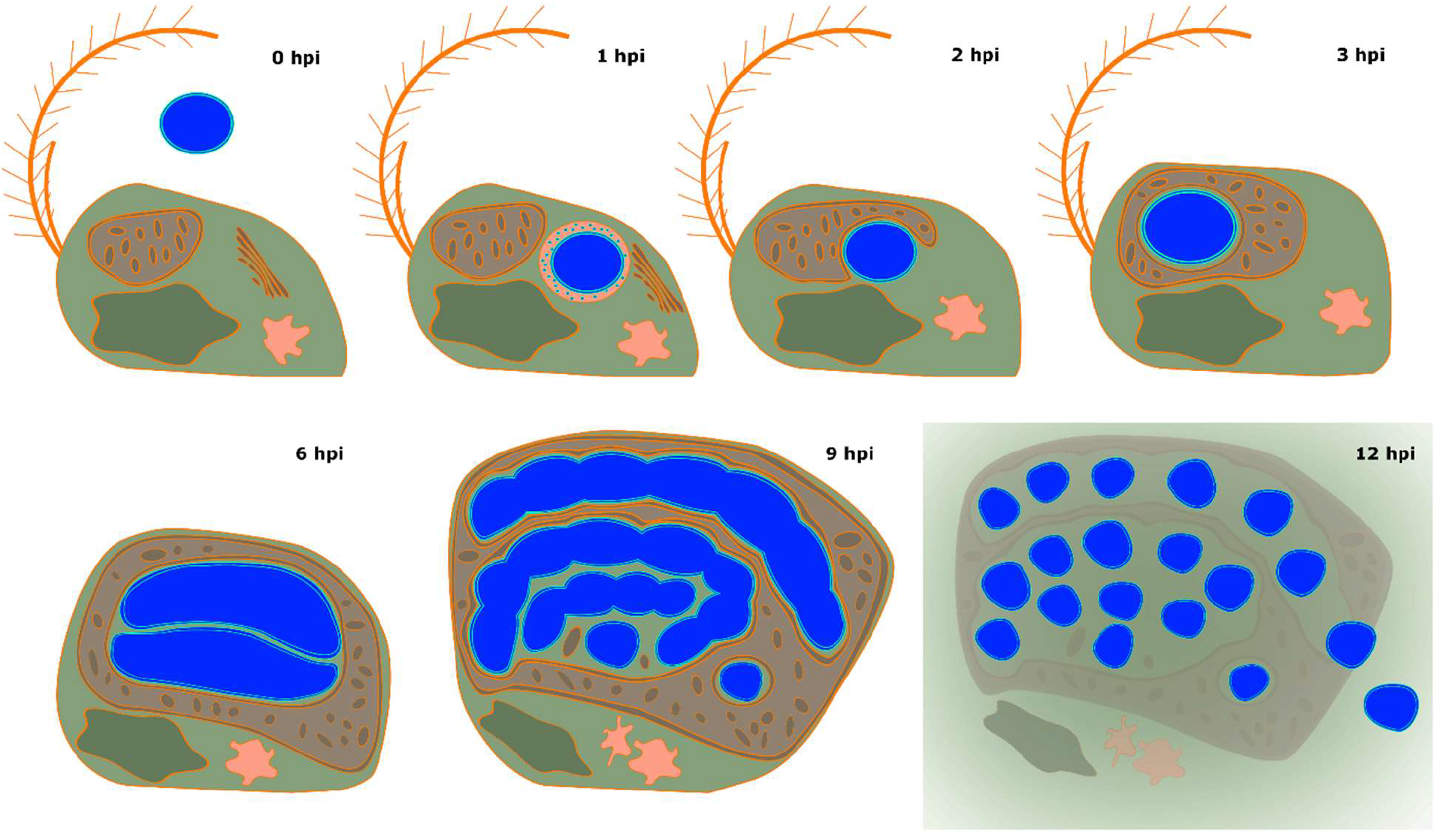
*Schematic representation of the inferred replication cycle of Chromulinavorax destructans*. The host S. elongata cell is shown in green, with the mitochondrion in brown, nucleus in dark green, Golgi in orange, and vacuoles in light orange. C. destructans is depicted in blue throughout its replication cycle.

### The *Chromulinavorax destructans* genome encodes a minimal set of core functions but an extensive assortment of genes involved in host modification

The 1,174,272 bp circular ds DNA *Chromulinavorax destructans* genome has a GC content of 33.7%, similar to other intracellular parasitoids of eukaryotes [6, 29]. GC skew and the location of the presumptive DnaA box ”TTATCCACA” suggest that the origin of replication lies at 651641-652233bp (Figure 6A). The genome encodes two typical complete rDNA loci and 35 tRNAs covering all twenty amino acids (Figure 6A). Other ncRNAs include 4.8S RNA (*ffs* ncRNA), RNase P RNA (*rnpB* ncRNA), as well as a tmRNA (*ssrA*) (Figure 6A). Of the 1,081 predicted open reading frames 55% had a functional prediction with the largest fractions involved in DNA replication, translation, trans-membrane transporters and host manipulation (Figure 6B).

**Figure 6.**
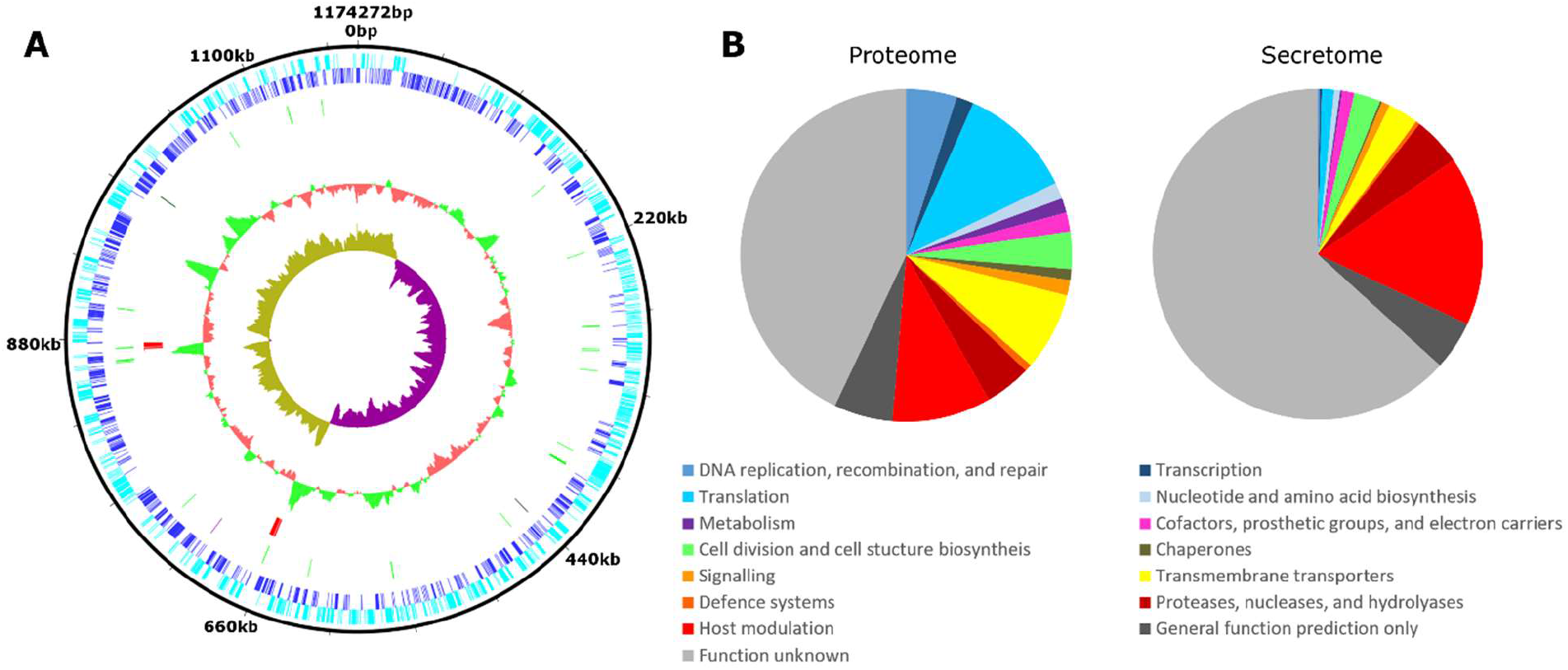
*Genome content of Chromulinavorax destructans*. A: Genome map showing open reading frames encoded by the plus and negative strand (dark and light blue respectively), tRNAs (green), rRNAs (red) and other ncRNAs (black). Inner circles depict GC content (red/green) and GC skew (teal/purple). B: Functional groups of putative protein-coding genes (Proteome) and the fraction coding a signal sequence (Secretome).

The DNA replication and repair machinery of *C. destructans* consists of a simple origin binding complex (lacking *dnaC*) and a typical prokaryotic replicase complex. DNA repair pathways include nucleotide excision repair (uvr system), short- and long-patch base-excision repair and mismatch repair (*mutL/S/Y* and *recJ*). Homologous recombination is supported by the *rec* and *ruv* pathways (with dpo), as well as site-specific recombinases (*xerJ/D*). The RNA polymerases assisted in transcription by several transcriptional regulators (*merR, yebC, fmdB*), transcription termination factors (*nusA/B/G*), and RNase H.

The ribosome, missing subunits 30S-22, 50S-8, 50S-25, 50S-26, 50S-30 and the rRNAs are edited by several ribosomal modification enzymes such as methyl and acetyl transferases. Similarly, the tRNAs are modified by a large number of modifying enzymes such as methyl transferases and dihydrouridine synthases and a full set of 21 aminoacyl-tRNA synthetases. Peptide chain release factors (*prfA/B*), ribonucleases (RNaseY/P), ribosome recycling factor (*rrf*), translation initiation and elongation factors promote translation (*infA/B/C, fusA, lepA, elf, tsf, tufB*). Post translational modifications are assisted by several chaperones and some proteins are subjected to transport to membranes.

*Chromulinavorax destructans* utilizes nucleoside salvage for both purine and pyrimidine biosynthesis and is able to phosphorylate the products into all required nucleotides. Therefore, precursor nucleosides must be imported from the host. Similarly, amino-acid biosynthesis is restricted to simple conversions between related amino acids (ser/cys, cys/ala, glu/gln, ser/gly), emphasizing that *C. destructans* is highly dependent on the host.

*Chromulinavorax destructans* encodes very rudimentary cell-division machinery, with only *ftsA, K, L, W*, and *Z* present, supported by *zapA*. Cell shape is determined by *mreB, C*, and *RodA*. Consistent with microscopic observations of a gram-negative-like phenotype, no LPS biosynthesis genes were observed. However, a complete *mur* pathway of peptidoglycan biosynthesis is likely responsible for generating the electron-dense material in the periplasm observed in electron micrographs (Figure 1D). Several surface antigens of unknown function are also encoded, as well as Type IV pilus assembly proteins, which might be assembled through a derived *pulD* channel. Several signaling trans-membrane receptors and signal-transduction proteins influencing the cell cycle are putatively involved, which could be involved in switching from dormancy to active replication.

As there was no evidence for complete metabolic pathways encoded by the genome of *C. destructans*, it implies that the cells must rely on extensive transmembrane transport systems to import metabolites and other resources from its host (Figure 7). A large array of ABC transporters involved in importing oligopeptides and amino acids through the *pot* and *opp/dpp* systems, presumably provide amino acids for protein synthesis. Phospholipids and lipoproteins are imported through the *mal* and *lol* ABC transporter systems respectively. ABC transporter systems *fep* and *tro* import trace elements such as iron, zinc and manganese (*znu*). Other ABC transporter systems of uncharacterized specificity are present, including putative multidrug exporters. Besides the ABC transporter systems, several specialized symporters, antiporters, and pumps are predicted to import potassium (*trk*), sodium (*nha/als/put*), inorganic ions (*mgt*), and nucleosides (*nup*). Further, specialized multidrug exporters are also present (*rhat/eama*). Central to energy requirements, there are two copies of a *tlcc* ATP/ADP antiporter that allows for the exchange of ADP for ATP from the host, which seems to be the only source of ATP. The membrane potential, essential for many transporters and antiporters, appears to be maintained by an ATP-synthase running “in reverse”, which may be regulated by passive proton channels. Large biomolecules can be imported by mechanosensitive channels (*msc*) and a biopolymer transporter (*exb*).

**Figure 7:**
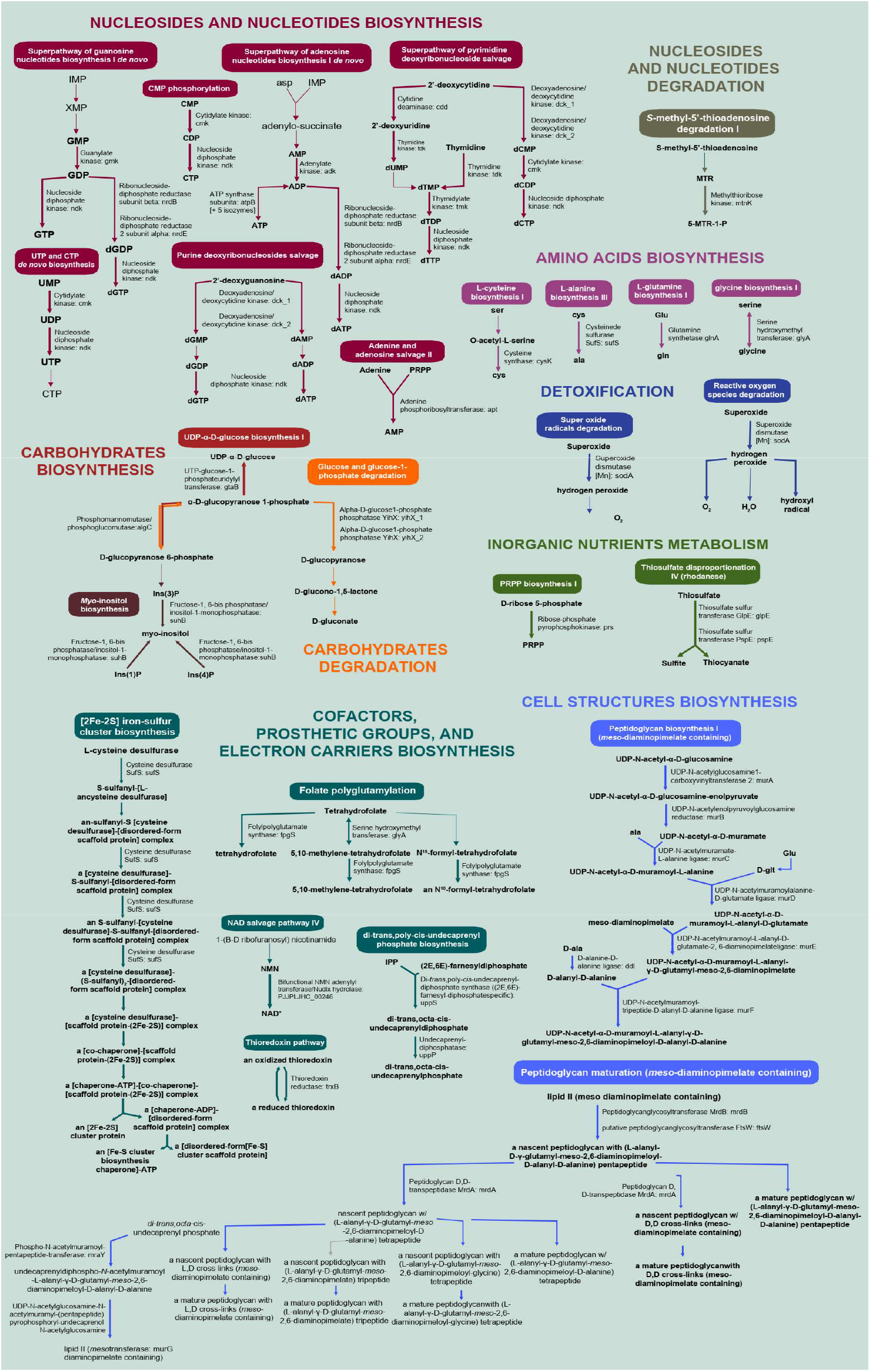
Chromulinavorax destructans metabolic pathways reconstructed by pathway tools

A substantial proportion of the genome encodes proteases, nucleases, and hydrolases, which are putatively involved in processing and breaking down host-cell structures and biopolymers that can be imported and reused. These effectors can be classified as secreted, membrane bound, and cytoplasmic factors. Amongst the secreted hydrolases is a close relative of chitinases and several HAD superfamily hydrolases (*yigB, mphC, glcD*). Secreted proteinases include M3, M15, M23, M41, M48, C11, and C65 peptidases. C65, M23, and M50 peptidases are predicted to be membrane bound. The final step in host biopolymer degradation is accomplished by cytoplasmic endonucleases (*endoU*), hydrolases (NUDIX and HIT-type), and proteases (lon, M16, M17). As well, a putative superoxide dismutase is encoded, which could help the cell cope with stressors such as hydrogen peroxide from close proximity to the mitochondrion.

More than 10% of the proteins encoded by *C. destructans* are predicted to be involved in modifying and influencing the host cell. Most prominent are 98 copies of ankyrin-repeat domain proteins. This concurs with observations of other intracellular parasitoids among the TM6 candidate phylum, such as *Babela massiliensis*, but also includes unrelated intracellular parasites such as *Legionella* spp., and giant viruses [6, 20, 30, 31]. The exact function of ankyrin-repeat domain proteins is unknown, but they have been implicated in membrane modification and counteracting the host immune system [30, 32, 33]. A CDS for a protein distantly resembling mitofilin was also found, and is an intriguing candidate for the extensive manipulation of the mitochondrion.

A *sec* secretion system is likely used to export proteins in the periplasm and to secrete proteins. Interestingly, 44% of all CDS possess a putative signal peptide that targets them to the secretion system, either as membrane proteins or as secreted proteins. This subset of secreted proteins is enriched in ORFans and proteins of unknown function compared to the complete set of CDS (Figure 6B). However, other overrepresented fractions include putative CDS for proteins that manipulate the host, such as proteinases and nucleases, and most prominently the ankyrin-repeat domain proteins, 78% of which contain a signal protein.

### Phylogenetic position of *C. destructans* within the candidate phylum TM6/Dependentiae

Full length 16S rDNA maximum likelihood (ML) phylogenetic analysis of the TM6/Dependentiae places *C. destructans* into the proposed *Babeliales* (ord. nov.), within the *Babeliae* (class nov.) [4], but within a sister family to the *Babeliacea* (fam. nov.), for which the name *Chromulinavoracea* (fam. nov.) is proposed. This family also contains the original MAG of JCVI_TM6SC1 and might also harbour *Vermiphilus pyriformis*, based on a partial 16S sequence [5]. The basal nodes in the 16S rDNA ML tree are poorly supported, likely due to undersampling, and similar to previous analysis (Figure 8A) [4]. The phylogenetic position of *C. destructans* was also explored by ML analysis of concatenated ribosomal proteins from MAGs that confidently contained all of the ribosomal CDSs investigated. The resulting tree reflects the architecture of the 16S rDNA tree and supports the proposed sister families *Babeliacea* and *Chromulinavoracea* within the order *Babeliales*, which is distinct from a second well-supported order of environmental sequences (Figure 8B). Genome content analysis of representatives of the two families within the *Babeliales* supports the separation of the families *Babeliacea* and *Chromulinavoracea*. The proposed members of the *Chromulinavoraceae* share more clusters of orthologous genes with each other than with *Babela masiliensis* of the *Babeliacea*, despite the *Chromulinavoracea* including potentially incomplete MAGs (UASB293, SOIL82, and JCVI TM6SCI) and the *Babeliacea* being represented by a complete genome (Figure 8C).

**Figure 8:**
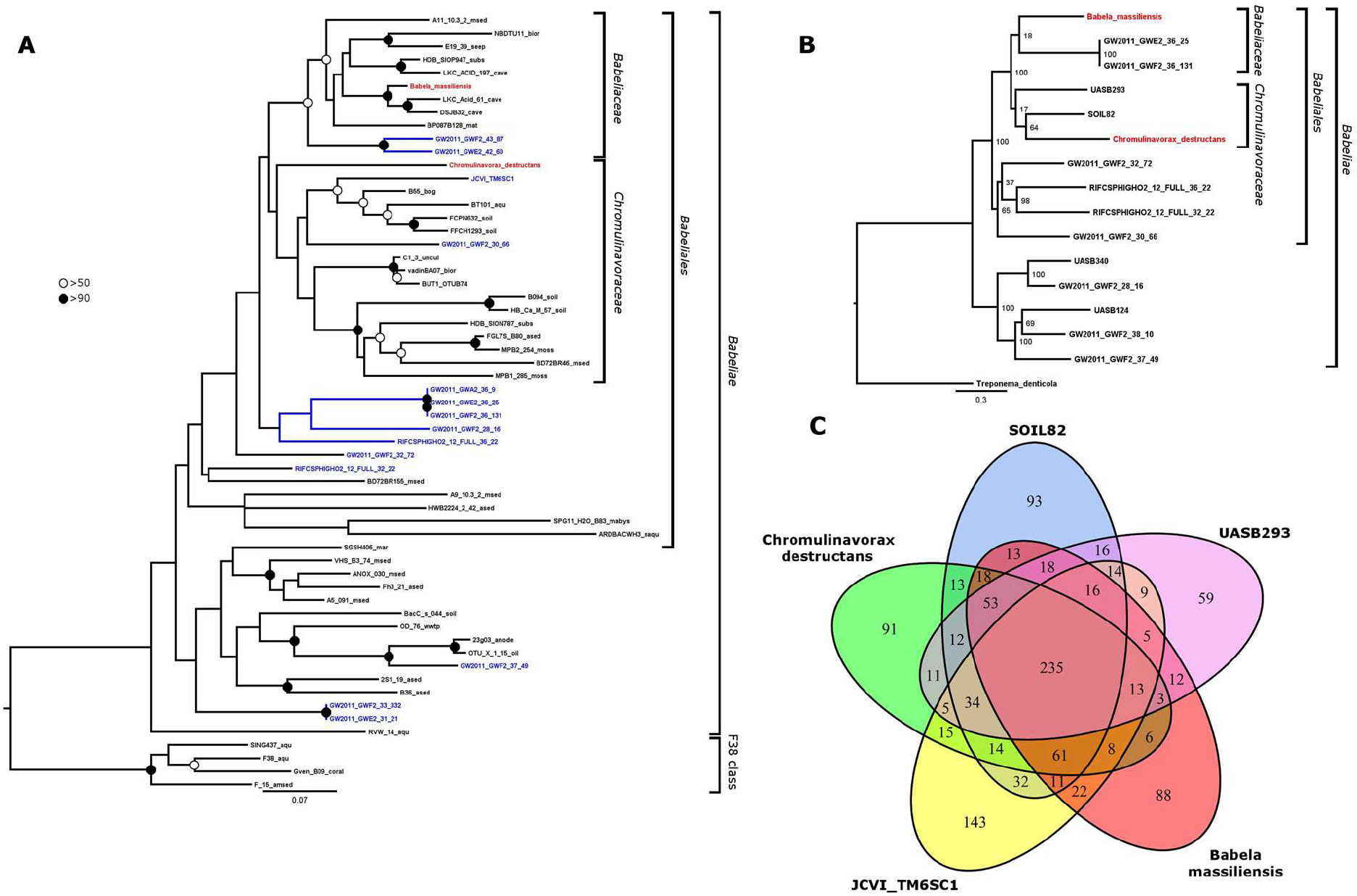
*Phylogenetic placement of Chromulinavorax destructans*. A) 16S full length rDNA ML tree (1000 BS replicates) showing the TM6/Dependentiae phylum. “Substantially complete” MAGs highlighted in blue, isolates in red. Solid circles show node support of >90 and open circle >50. B) ML tree of ribosomal proteins (L2, L3, L4, L5, L6, L14, L15, L16, L18, L22, L24, S3, S8, S10, S17, S19; BS support based on 1000BS replicates). C) Shared gene clusters between *C. destructans* and members of its family *Chromulinavoracea* (UASB293, SOIL82, and JCVI TM6SCI) and *Babela massiliensis* as a representative of the *Babeliacea*.

## Discussion

### Chromulinavorax destructans is highly host-dependent

*Chromulinavorax destructans* is an intracellular pathogen of *S. elongata* that appears to be dependent on its host for replication, to the extent that it does not appear to encode any complete metabolic pathways (Figure 6,Figure 7). Many putative effector molecules, such as proteases, nucleases and ankyrin–repeat domain proteins, that could manipulate and break down host structures contain signal peptides and appear to be secreted from the bacteria (Figure 4A). Similarly, putative outer membrane vesicles that could also contain such effector molecules bud during the early stages of infection and might serve a similar function (Figure 4A). The reorganization and breakdown of host structures provide resources, possibly due to aforementioned effector molecules, that can be imported into the C. *destructans* replication bodies (Figure 9). With no evidence of lipid biosynthesis, in combination with the observed host membrane disarray during late infection, suggests that host lipids are used for the bacterial cell membrane. Remodeling of the host mitochondrion is one of the most unusual features of the replication process. The expansion of the mitochondrion while maintaining membrane integrity contrasts with observations from other mitochondrion-invading pathogens and symbionts that actively disrupt the mitochondrial integrity [34, 35]. The proximity of bacterial replication to the expanded mitochondrion would allow for a steady supply of ATP mediated by the encoded ADP/ATP antiporter, but it would also require free radicals to be neutralized, a function likely performed by bacterial encoded superoxide dismutase. This high host-dependence is consistent with the pathogen being dormant until taken up by a host cell, and the lack of evidence of genome replication in free bacteria. It may also be the reason that the cells remained infectious after four years at 4°C. Complete host dependence and extracellular dormancy are life-cycle characteristics that highlight traits of convergent evolution found in many obligate pathogens, including viruses and some of the most reduced eukaryotic parasites. For example, Microsporidia completely lack energy generation pathways, while ATP transporters are widespread [36–38]. The high similarity in life style and genome content between TM6/Dependentiae and giant viruses, provides an intriguing example of converging evolutionary trajectories from vastly different evolutionary backgrounds.

**Figure 9:**
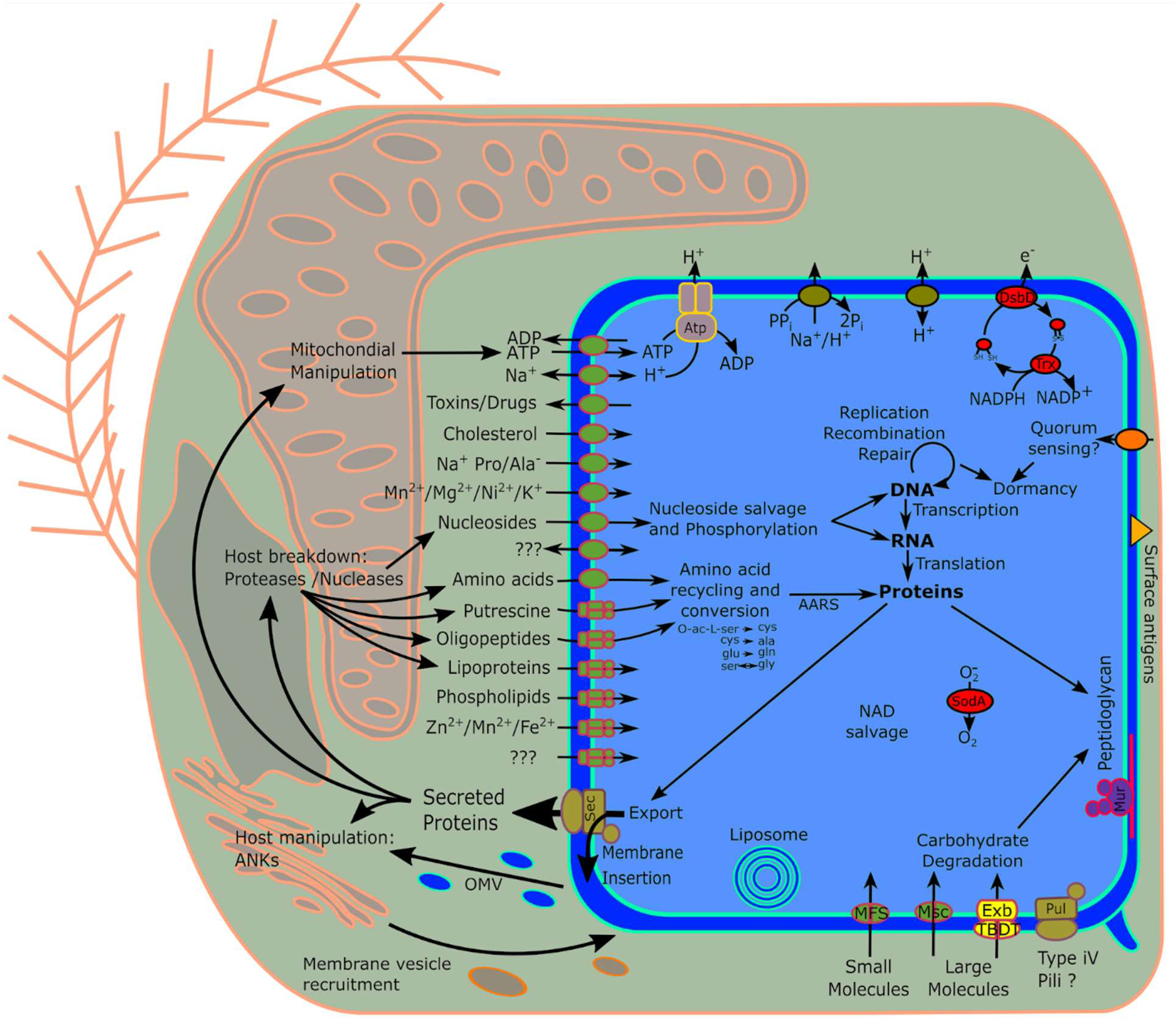
Schematic representation of inferred metabolic capabilities of Chromulinavorax destructans and its interactions with its host, Spumella elongata.

### The extensive mitochondrial modifications caused by *Chromulinavorax destructans* might stem from limited host resources

Rudimentary cell-division machinery and the lack of many hallmark genes has been highlighted for the TM6/Dependentia group, specifically based on MAGs [3, 4]. Similarly, *Babela massiliensis* showed delayed cell division after initial growth in amorphous bodies (i.e. replication bodies), and, while not directly described, *Vermiphilus pyriformis* likely uses a similar replication strategy [5, 6]. Although replication of *C. destructans* in replication bodies is much like *B. massiliensis*, the close association and modulation of the mitochondrion was unique to *C. destructans*. In part, this difference might reflect the availability of cytoplasmic resources, such as metabolites and ATP, that presumable are more scarce in the flagellate cell compared to the much larger amoeba; Accordingly, *C. destructans* causes severe deformation of the cellular architecture of the nanoflagellate that was not seen in amoeba hosts (Figure 3, Figure 4). The scarcity of cytoplasmic ATP could lead to the close association of *C. destructans* with the host mitochondrion and initiate the mitochondrion’s expansion and rearrangement. A limitation of energy could have led to the incomplete replication bodies that were frequently found inside ghost cells, suggesting that ATP availability might restrain *C. destructans* replication. The differences between amoeba-infecting and nanoflagellate-infecting TM6/Dependentiae resemble observations made for giant viruses infecting different hosts. Nanoflagellate-infecting giant viruses show highly spatially oriented replication and preserve the mitochondrion; whereas, amoeba-infecting viruses seem to be unrestricted by the host-cell architecture [20, 39]. Additionally, a similarly high percentage of the genome of the lytic TM6/Dependentiae representatives is dedicated to ankyrin-repeat domain proteins, suggesting that these are crucial effectors for intra-eukaryotic replication and host cell take-over [6, 20]. Although their specific function of these proteins within these host-pathogen systems remain unknown, their nature as protein-proteins interactions domains suggests they might be involved in host modification strategies [30]. Where ankyrin-repeat domain proteins have been explored in other host-pathogen systems, their functions include blocking of the host immune system, the induction of phagocytosis, the facilitation of cytoplasmic invasion, and the self-protection from potentially damaging effector molecules [30, 33, 40–42]. Any one of these functions could benefit *C. destructans* during its replication and therefore might be deployed.

### Life history strategies and host-range might differ vastly amongst the TM6/Dependentiae and might promote diversity within the phylum

Phylogenetic analysis confidently places *C. destructans* within the candidate phylum TM6/Dependentiae, making it the first reported isolate of the phylum that does not infect an amoeba, and the first to infect a representative of the stramenopile supergroup of eukaryotes. Other studies have reported a correlation of TM6/Dependentiae with ciliates, suggesting that they infect or are symbionts of other eukaryotic lineages [4]. Life-history strategy seems to be independent of the host taxonomic group, given that *C. destructans* and *B. massiliensis* are lytic, but infect distantly related hosts, while *V. pyriformis* is symbiotic, and like *B. massiliensis* infects an amoeba. This suggests that life-history strategy, and resulting pathogenicity, is a product of coevolution within a specific host-pathogen pair. The restricted host ranges of *C. destructans* and *V. pyriformis* are consistent with members of the TM6/Dependentiae being highly adapted to their hosts. The highly restricted host range of members of the TM6/Dependentiae suggests that they are extraordinarily diverse, as is reflected by the great diversity of related 16S rRNA sequences from environmental surveys. Given their potentially broad host range and distribution, TM6/Dependentiae could play a significant role in controlling the abundance and diversity of aquatic autotrophic and heterotrophic protists.

### Summary

*Chromulinavorax destructans* is the first described isolate of the candidate phylum TM6/Dependentiae that does not infect *Acanthamoeba* and is the first isolate of a novel family so far only represented by MAGs. It causes extensive reorganization of the host cell, most notably the mitochondrion wrapping around the bacterium’s replication bodies. The genomic complement of *C. destructans* is highly reduced in metabolic capabilities, encoding not a single complete metabolic pathway and instead relies extensively on host resources such as metabolites and even ATP for energy supply. In contrast, the *C. destructans* is rich in putative effector molecules, putatively breaking down and reorganizing the host cells, as well as a large number of uncharacterized proteins and proteins not represented in databases. *C. destructans’* narrow host range provides hints to a regulatory role of parasitic bacteria in the diversity and abundance of heterotrophic protists

## Acknowledgements

The work was supported by grants to C. S. from the Natural Sciences and Engineering Research Council of Canada (NSERC; 05896), Canada Foundation for Innovation (25412), British Columbia Knowledge Development Fund, and the Canadian Institute for Advanced Research (IMB). C. D. was supported in part by a fellowship from the German Academic Exchange Service (DAAD). F.H. is supported by a postoctoral fellowship from EMBO (ALTF 1260-2016)

